# CellSpecks: A Software for Automated Detection and Analysis of Calcium Channels in Live Cells

**DOI:** 10.1101/355693

**Authors:** S I Shah, M Smith, D Swaminathan, I Parker, G Ullah, A Demuro

## Abstract

To couple the fidelity of patch-clamp recording with a more high-throughput screening capability, we pioneered a novel approach to single channel recording that we named “optical patch clamp”. By using highly-sensitive fluorescent Ca^2+^ indicator dyes in conjunction with total internal fluorescence microscopy techniques, we monitor Ca^2+^ flux through individual Ca^2+^-permeable channels. This approach provides information about channel gating analogous to patch-clamp recording at time resolution of ~ 2 ms, with the additional advantage of being massively parallel, providing simultaneous and independent recording from thousands of channels in native environment. However, manual analysis of the data generated by this technique presents severe challenges as a video recording can include many thousands of frames. To overcome this bottleneck, we developed an image processing and analysis framework called CellSpecks, capable of detecting and fully analyzing the kinetics of ion channels within a video sequence. By using a randomly generated synthetic data, we tested the ability of CellSpecks to rapidly and efficiently detect and analyze the activity of thousands of ion channels, including openings for a few milliseconds. Here, we report the use of CellSpecks for the analysis of experimental data acquired by imaging muscle nicotinic acetylcholine receptors and the Alzheimer’s disease-associated amyloid beta pores with multiconductance levels in the plasma membrane of *Xenopus laevis* oocytes. We show that CellSpecks can accurately and efficiently generate location maps, create raw and processed fluorescence time-traces, histograms of mean open times, mean close times, open probabilities, durations, and maximum amplitudes, and a ‘channel chip’ showing the activity of all channels as a function of time. Although we specifically illustrate the application of CellSpecks for analyzing data from Ca^2+^ channels, it can be easily customized to analyze other spatially and temporally localized signals.

## INTRODUCTION

Ca^2+^ is a highly specific universal second messenger that regulates numerous cellular functions and plays a key role in many diseases [1–8]. In physiological conditions, Ca^2+^ is tightly regulated by ion channels, pumps, and buffering proteins [8–10]. Thanks to the recent progress in imaging technology and the development of more efficient Ca^2+^-sensitive dyes, it is now possible to image Ca^2+^ signals inside intact cells with high spatiotemporal resolution, reaching the ability to monitor Ca^2+^ flow across single plasma membrane Ca^2+^-permeable channels including, voltage-sensitive N-type Ca^2+^ channels [11,12,14,15], L-type Ca^2+^ channels in cardiac muscle, dihydropyridine-sensitive voltage-gated Ca_V_1.2 channels [17], ligand-gated muscle nicotinic acetylcholine receptors (nAChRs) [18], inositol 1,4,5-trisphosphate (IP_3_) receptors (IP_3_Rs) [16], and membrane pores formed by soluble amyloid beta oligomers (Aβ42) characterized by multiple conductance levels [11–17]. Surprisingly, a common feature of all these channels is their lack of motility as the florescent signal generated at a specific site can be imaged for many tens of second suggesting a stationary feature during channel activity [12,14].

We have pioneered an imaging technique called ‘optical patch clamp’, a massively parallel 2D optical approach, capable of simultaneously and independently monitoring the functions of several hundred Ca^2+^-permeable channels at the single channel resolution [18]. Binding of Ca^2+^ to cytosolic fluorescent Ca^2+^ indicators, generates fluorescent signals (Single Channel Calcium Fluorescent Transients; SCCaFTs) that closely track the opening and closing of ion channels [15]. The presence of these bright spots on the 2D video sequence, captured by total internal reflection fluorescence microscope (TIRFM), can be identified manually by visual inspection to infer channel location and activity. However, the manual detection of these elementary events is extremely tedious and time consuming, and does not lend itself to a comprehensive analyses of the stacks as it is nearly impossible to accurately determine the location of each of the hundreds or sometimes thousands of channels present in the image field. This is particularly difficult for channels with extremely low open probability (P_o_) and short opening events where the few recorded events are several thousand frames apart. This could result in an incomplete representation of the channel population behavior.

Several automated detection packages are available for analysis of similar data, such as GMimPro for single fluorophores [19]; ImageJ plugins for line-scan images like SparkMaster [20] and xySpark [21]; FLIKA for Ca^2+^ puffs resulting from concerted gating of clustered IP_3_Rs [22]; and several algorithms for processing of Ca^2+^ sparks generated by clustered ryanodine receptors (RyRs) [23–29]. However, the existing software packages are limited in their abilities to detect a large number of simultaneously active channels (sites) with multiple conductance levels and generate huge data-sets about the location and gating kinetics of these channels. For example, in our experiments, typical imaged data consists of a high temporal resolution image stack recorded at ~500 frames per second, in a 128 pixel × 128 pixel multiimage (up to 15,000 frames) Metamorph (.stk) file. Automated detection of channel locations and their gating kinetics would allow for a through description of the behavior of every channel, reliably pinpointing all event locations regardless of size, frequency, and duration. Furthermore, statistical analysis can be customized and automated, providing a powerful tool for ion channels and single molecule studies.

With these needs in mind, we developed an automatic detection algorithm, coupled with graphical user interface and statistics modules. We named the resulting software CellSpecks, which is implemented in the Java programming language for speed, flexibility, and portability. CellSpecks is capable of automatic detection of the location and gating behavior of many ion channels in Metamorph Stack file or image libraries in formats such as JPG and TIFF, with tremendous efficiency over manual approach and able to reliably identify subtle events where manual identification is tedious if not impossible. As a result, analysis of imaging data from TIRFM experiments, now takes only a fraction of the time needed for visual inspection (only a few minutes instead of many hours and days), and events previously not accessible manually or through other packages are made available for statistical analysis.

Here, we first use CellSpecks to analyze synthetic images sequences where all events are known a priori to validate its accuracy in a challenging but anticipated situations. After validating the performance and accuracy, we then illustrate the use of CellSpecks to process TIRFM image sequences of Ca^2+^-permeable nAChRs channels and individual pores formed by Aβ42 oligomers in the plasma membrane of *Xenopus laevis* oocytes. CellSpecks can efficiently generate location maps, create raw and processed fluorescence time-traces, histograms of maximum amplitudes of Ca^2+^ release events, mean open times, mean close times, and P_O_ of all channels, and amplitudes and durations of all events in the image sequence. All data can be exported in ascii format for plotting and further analysis. In particular, time-traces from all channels can be exported and idealized using the open source software TraceSpecks [30, 31] for developing single channel models. Thus, CellSpecks together with the high-resolution fluorescence microscopy provides a powerful tool for characterizing and modeling ion channels behavior, using unprecedented amount of data sets simultaneously recorded from thousands of channels in their native environment.

## MATERIALS AND METHODS

The experimental data used to test CellSpecks derive from previously published work imaging the activity of nAChRs [14] and pores formed by Aβ42 oligomers [13]. A brief description of these methods is given below.

### Oocyte preparation and electrophysiology

Experiments were performed on defolliculated stage VI oocytes obtained from *Xenopus laevis* [32]. For experiments with muscle nAChRs, in vitro-transcribed cRNAs coding for α, β, γ and δ subunits (in a molar ratio 2:1:1:1) were mixed to a final concentration of 0.1-1 μg/μl and microinjected (50 nl) into oocytes [18]. After 3-5 d the expression of nAChRs was monitored by recording currents evoked by bath application of ACh [18]. Insertion of functional Aβ42 pores into the oocyte’s plasma membrane [13], was achieved by bath application of solution containing soluble oligomers prepared from human recombinant Aβ42 peptide. The solutions containing Aβ42-oligomers were delivered from a glass pipette with tip diameter ~ 30 μm positioned near the edge of the membrane footprint of the oocyte membrane on the cover-glass.

Oocytes were injected ~ 1 hr before imaging with fluo-4-dextran (MW ~10 kD; Ca^2+^ affinity about 3 μM) to a final intracellular concentration of ~40 μM. For imaging, oocytes were placed animal hemisphere down in a chamber whose bottom is formed by a fresh ethanol washed microscope cover glass (type-545-M; Thermo Fisher Scientific) and were bathed in Ringer’s solution (110 mM NaCl, 1.8 mM CaCl_2_, 2 mM KCl, and 5 mM Hepes, pH 7.2) at room temperature (~23°C) continually exchanged at a rate of ~ 0.5 ml/min by a gravity-fed superfusion system. The membrane potential was clamped at a holding potential of 0 mV using a two-electrode voltage clamp (Gene Clamp 500; Molecular Devices) and was stepped to more negative potentials-100 mV when imaging Ca^2+^ flux through the gating channels to increase the driving force for Ca^2+^ entry into the cytosol [13, 18].

### TIRFM imaging

Imaging was accomplished by using a custom-built TIRF microscope system based around an Olympus IX 71 microscope equipped with an Olympus 60x TIRFM objective (NA = 1.45) [13,18]. Fluorescence excited by a 488 nm laser was imaged using an electron-multiplied ccd camera (Cascade 128+: Roper Scientific) at full resolution (128 × 128 pixel: 1 pixel = 0.33 μm at the specimen) at a rate of 500 s^−1^. Image data were acquired using the MetaMorph software package (Universal Imaging, Westchester, PA).

### Event detection algorithm and automated analysis

CellSpecks has a menu-driven interface for reading and processing video stacks and individual images, as well as the ability to illustrate various steps and results of the algorithm, allowing the experimenter to confirm the effectiveness of the algorithm for their particular dataset. Third party libraries capable of returning pixel byte-arrays for these video/image stacks are used to feed image data into the program that convert an image stack from its native format to a stack data-structure. The stack is then cloned (copied) and modified a number of times along the way in order to preserve processed data that will later help the experimenter understand how the algorithm produced the results.

Figure 1 summarizes the steps, including noise detection, signal isolation from noise, event attribution, and analysis from time-trace of a given channel. After cloning the initial input movie frames or tiff images, the program takes each pixel in the image over time and assigns it to an individual thread to generate blurred signal, noise, and signal stacks from the **Original** stack. Flowchart 1 in Supplementary Information Text 1 lists various steps involved in computing **Noise** stack from **Original** stack. These steps are explained below.

**Figure 1.**
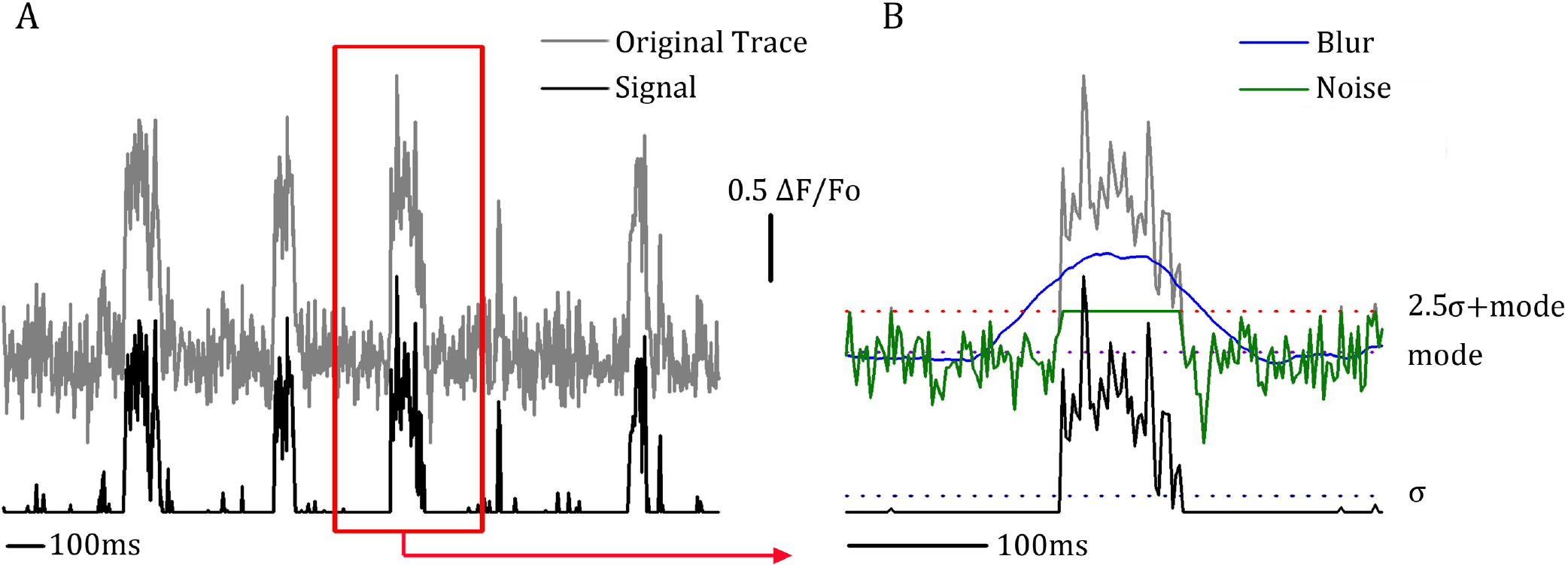
Illustration of CellSpecks algorithm processing steps - separation of noise from signal nAChRs channel activity. (A) shows fluorescence trace from a single pixel (gray) and the final signal trace (black) calculated by CellSpecks. (B) Illustrates processing steps required to go from the original (gray) trace to the signal (black) trace. First temporal smoothing creates a blur (blue) trace. A mode value is calculated for the entire blur trace (purple dotted line). Standard deviation, σ, is calculated for all fluorescence points from the original trace that are below the mode value (dark blue dotted line). Noise threshold is set as 2σ+mode (red dotted line) and a noise trace generated (green). Subtracting noise (green trace) from the original trace (gray) the signal trace (black) is obtained. In this figure all time varying quantities (original trace, blur, noise, signal) are plotted with solid lines whereas time-invariant variables (mode, σ, noise threshold) are plotted with dotted lines.

First, the **Original** stack is copied to a temporary stack called **Temp**. Each pixel in the **Temp** stack is then replaced by the average of the intensities of the same pixel over time (51 frames) by the application of a one-dimensional flat kernel of length 51 (a moving box filter with a window size of 51 frames) with the center of the kernel being the intensity value to be modified. Appropriate boundary conditions are applied at the boundary (both in time and space) pixels. As a result, a modified stack (**Blur**) is generated for each pixel. Next, a mode value for each pixel over time in the **Blur** stack is determined. This is followed by computing the standard deviation of the points in the **Original** signal that are below the mode of this pixel. A noise threshold is set at 2.5 × standard deviation + mode. All intensity values over time of this pixel in the **Original** stack which are greater than the noise threshold are replaced with the mode value to get what is called **Noise** stack. Subtracting **Noise** stack from the **Original** image stack generates a **Signal** stack. The **Signal** stack is then used to identify all Ca^2+^ channels along with their location, open, close events, PO, and maximum amplitudes. Various steps involved in event detection, channel locations, and assigning events to different channels are listed in Flowchart 2 in Supplementary Information Text 1 and described below.

First a three-dimensional array of size X×Y×Time (where X and Y are the dimensions (number of pixels along x and y axes) of the image and Time is the number of images(frames) in the data), called *EventPartStack* is created to hold each pixel’s data (intensity) over time. Each element of this array also contains variables to link each non-zero intensity pixel (pixel with no signal is represented by a value of zero) to four neighboring pixels in the current frame (at time t) and two neighbors in the frames before (time t-1) and after (time t+1) the current frame. Each non-zero element of the *EventPartStack* array is linked to all of its non-zero six neighbors. Now for each non-zero value of *EventPartStack*, an event, containing links to the non-zero neighbors of current pixel are recursively generated unless a zero-valued pixel is reached, and added to probable event list (*probEventlist*) array. The size of the*probEventlist* array now contains all probable events for all the channels in the given data set. This *probEventlist* is further screened for events whose duration is < 10 frames or whose cumulative intensity is < 50 (intensity units) and a final list of events called *Eventlist* is generated.

In order to accurately place these channels, a weight array is generated as a two-dimensional array (of image size) containing the sum of pixel values in the **Signal** over time for that pixel. The weight array is then normalized by the highest pixel value in the weight array. Non-zero values in weight array are interpreted as likely channel locations and added to a list of channels with no events.

The events are then added to channels one by one, either to the closest known channel if there exists one with which it overlaps, or to a new channel if no current channel overlaps. Channels are allowed to have floating point coordinates, complicating somewhat the concept of overlapping, especially when each event covers an area often no more than two pixels across. The nearest neighboring channel of each event is determined either by matching the coordinates of the event and channel or by calculating the overlap likelihood in case the coordinates do not have a one-to-one match (see “Channel Locations” section of Supplementary Information Text 1). The value of overlap likelihood is a combination of the inverse of the distance and the relative intensity of the event at the location of the nearest-neighbor channel. Thus, if the center of the channel does not overlap the event at all the overlap likelihood is zero, and any positive overlap value decreases to zero as the distance increases. Finally, channels containing no events are removed. At the end of this process, channel locations are adjusted for the actual events they contain and any channels found to be within the same pixel are combined. Further details about assigning events to channels and identifying channel locations are given in the “Channel Locations” section of Supplementary Information Text 1.

By the end of the detection and association process a list of channel locations has been produced, and within each channel is a list of known events, and in each event, is a list of known event parts (start and end of an event) as shown in Figure 2. The times between these events are averaged to give a mean close time, and durations of these events to give a mean open time. The total open time divided by the total sample time gives the P_O_. Maximum amplitude is the highest peak in the channel after a 1D smoothing kernel of length three (moving box average with a window size 3) is applied to the intensities over time to filter high frequency noise. The program also saves the durations and amplitude of all events of all channels in the stack or image sequence. Descriptive statistics as well as intermediate stack, baseline, signal, and noise information can be viewed in the CellSpecks graphical user interface (Figure 2). Channel locations are visually reported for one-click, intensity versus time graph confirmation, and the sampling for these traces can be modified to select an average or maximum of pixel values over a user-specified channel. These channel locations may also be threshold by maximum intensity. Statistical information, time-trace data (as modified by user-selected sampling parameters), location maps, and channel-chips (a surface (raster) plot showing the gating of all channels over time) may then be exported using the interface as well.

The resulting program has been tested on Mac OS X, Ubuntu Linux, and Windows XP-7, and using the Sun Java 1.6 JRE, Apple Java 1.5 and 1.6 JRE, and SoyLatte Java 1.6 JRE, and the OpenJDK Java 1.6 JRE. The pre-compiled JAR file, User Manual, sample stack file, and sample images are included with this paper as Supplementary Information Text 2. The source code for the software is available upon request from the authors. In trial runs against previously examined stacks, CellSpecks is capable of processing a 43pixel × 43pixel × 15,000 image stack in approximately 43 seconds on a 1.73GHz Intel Core i7-820QM with 8GB DDR3 RAM. The peak RAM usage during the detection process is ~7.5GB on Windows 7. This includes opening the images, feeding them into integer arrays, and executing the detection algorithms. Figure 2 shows typical confirmation images produced by the program.

**Figure 2.**
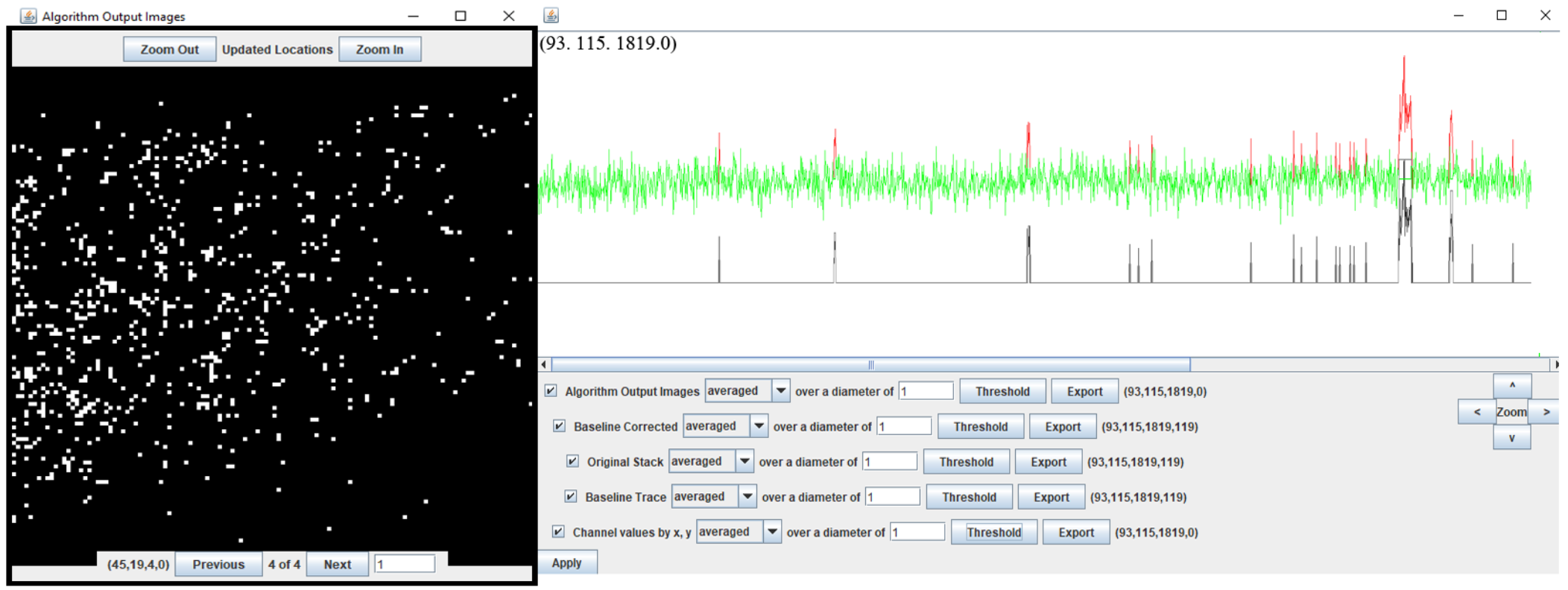
CellSpecks graphical user interface. CellSpecks interface includes the capacity to audit the detection and analysis process for quality and accuracy. The map of detected channels is shown in the Algorithm Output Images windows (left). Map is generated from a 5000 frames stack captured by imaging a 40×40 μm^2^ region of an oocyte plasma membrane expressing muscle (αβγδ) nAChRs. Channels are represented by bright pixels at their determined location. Brightness of a pixel corresponds to the maximum amplitude event generated by the channel. Clicking on any pixel brings up the trace window (right), displaying the time traces such as raw (red), baseline or noise (green), and processed or signal traces (black) of the channel at that location. Results of any modification are displayed in the Algorithm Output Images. Information such as mean open and close times, P_O_, peak amplitudes, life-times and amplitudes of all events for all channels, channel-chips showing the activity of all channels detected as a function of time, and channel location maps for all channels can be exported as ascii files for publication-quality plots and further analysis.

## RESULTS AND ALGORITHM TESTING

In the first part of this section, we use the time-traces, locations, and various statistics from all channels in the image sequence to verify that the output from the algorithm matches the ground truth of synthetic data.

### Synthetic data validation

To validate the accuracy of the algorithm, we generated a synthetic data set of 50 channels randomly distributed on a grid of 128 × 128 pixels for a total duration of 2 seconds in a sequence of 1000 tiff images. We deliberately randomized the opening and closing of these channels as well as their open and close times to mimic experimental conditions. To keep channel flux tractable, an amplitude of zero (intensity units) represents a closed channel whereas an open channel was assigned an amplitude of 200 (intensity units) in line with the frequently observed intensity values when the channel is open. To make the channel signal realistic, noise derived from normal distribution with mean of 5 and standard deviation of 2 was superimposed on each channel’s time-trace as well as all other pixels in the image frame. The observed spread of Ca^2+^ from the channel to the surrounding area is mimicked by allowing the fluorescence from an open channel’s location to spread to the nearest neighbor pixels. Code used to generate synthetic data is available on request from the authors. Further details about the algorithm generating the synthetic data are given in the Supplementary Information Text 1.

We have tested our algorithm for signal to noise ratios in the range of 5 to 40. CellSpecks was able to correctly identify channel locations, number of channels, and all opening events along with their open and close times as well as mean open and close times and Po for all signal to noise ratios (SNR) > 5. Figure 3 compares the results generated by CellSpecks for synthetic data set with the actual statistics of the data. As is evident from Figure 3A and 3B, the distributions for mean open and close times predicted by CellSpecks (second row) are in close agreement with the true distributions (top row). Similarly, the open probabilities estimated by CellSpecks (bottom) are in close agreement with the true values (top) (Figure 3C). The slight discrepancy in P_O_ distribution is due to the fact that CellSpecks excludes the partial open or close event towards the end of the recording, whereas such events were included in the true statistics. The gating state of all channels at a given time step (frame number) given by CellSpecks is saved as a channel chip representation (Figure 3D). The kinetics of all 50 channels given by CellSpecks are remarkably close to the true values. An example of synthetic channel trace generated with a SNR of 5 is shown in Figure 3E, whereas the time-trace identified by CellSpecks is shown in Figure 3F. A comparison of the two traces shows that CellSpecks is capable of identifying highly noisy channels along with open/close events precisely.

Although CellSpecks can accurately identify channels and their associated events for SNR > 5, for data with a SNR ≤ 5, we found that CellSpecks misses some channels as well as associated events. Figure 3G shows a comparison of the number of channels (left) and all events (right) identified by CellSpecks (green bars) with the actual values (red bars) as a function of SNR in the records. CellSpecks missed two channels for SNR = 5 but was able to identify all channels for SNR > 5. Similarly, CellSpecks missed 28 events for SNR = 5 while identified all events in images with higher SNR. Figure 3H shows a comparison between channel locations identified by CellSpecks (red circles for SNR = 10 and green circles for SNR = 5) and actual locations (x for SNR = 10 and yellow circles for SNR = 5). As is evident, channel location map identified by CellSpecks for SNR = 10 matches exactly with the actual locations. Same is the case for SNR = 5 except that CellSpecks missed two channels. It is worth mentioning that the SNR in TIRFM using fluorescence dye Fluo-4 is close to 8 and higher for Cal-520 [33]. CellSpecks did not detect any event in control stacks with no events.

**Figure 3.**
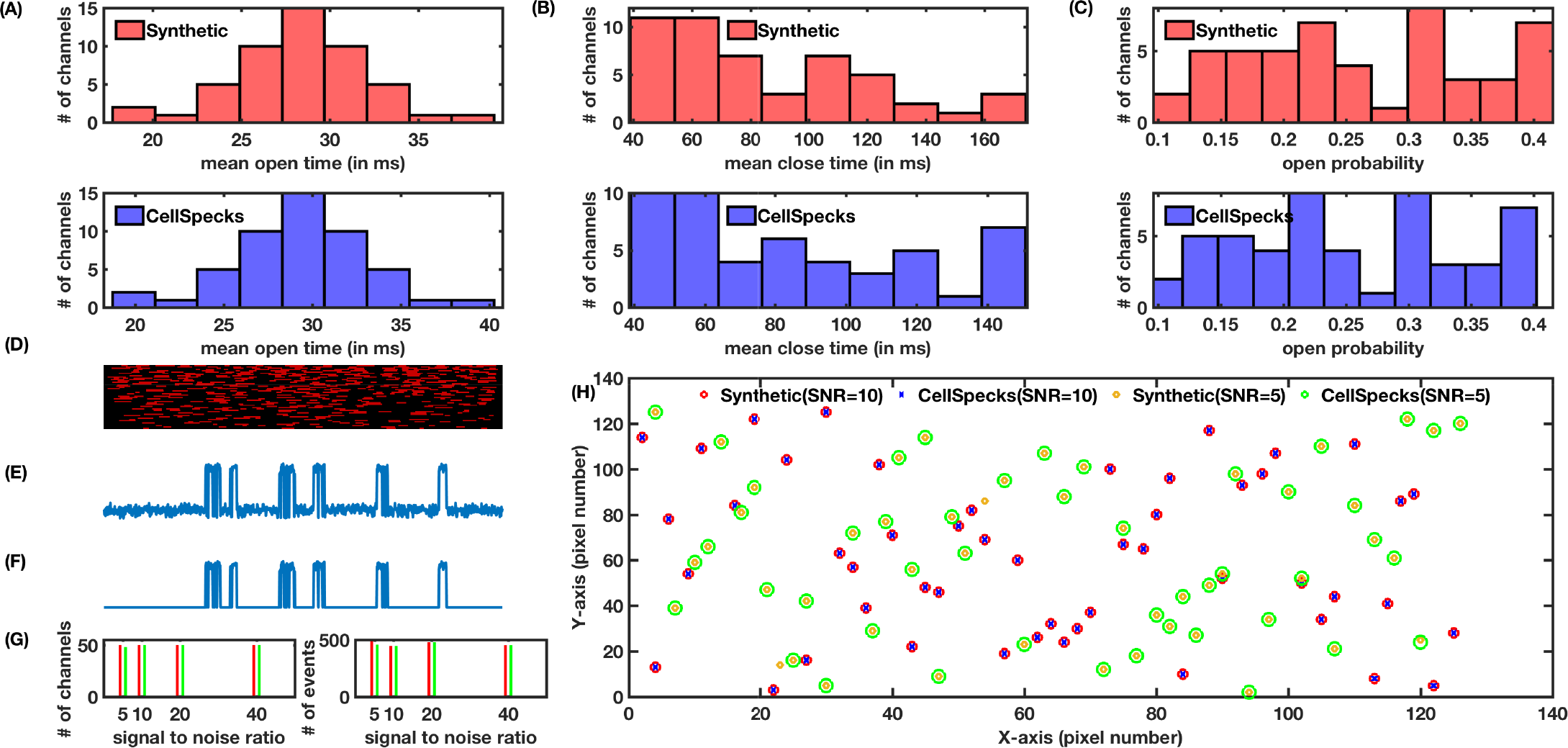
CellSpecks closely reproduces the channel locations and gating kinetics of simulated functional channels with random location and gating kinetics. Distributions of mean open times (A), mean closed times (B), and mean P_O_ (C) of true values (first row) and values estimated by CellSpecks (second row) for all channels. (D) Channel chip (image exported directly from CellSpecks) representation where horizontal axis is time (two seconds total) and vertical axis is the channel number (1 to 50), i.e. each horizontal line represents one channel. The red and black represent the open and closed state of the channel respectively. (E) Actual noisy (SNR = 5) time trace for a single channel and (F) the trace identified by CellSpecks. (G) The number of channels (left panel) and number of events (right panel) identified by CellSpecks (green bars) versus the actual values (red bars) as functions of SNR ratio. (H) Channel location maps from CellSpecks (green circles for SNR = 10 and red circles for SNR = 5) and true locations (x for SNR = 5 and yellow circles for SNR = 10).

We remark that event identification is accurate for all but extremely long events lasting more than 50% of the total duration of the recording. In cases where event duration approaches half of the total duration of the experiment, the amplitude of the event can be mistakenly considered to be the baseline. Note that the intensity during the event will have to be almost constant for this failure to happen. While unlikely for data without potentially compromising flaws, it could be an issue in data sets that have been cropped to represent a very short total duration.

### Detection of muscle nAChR channel activity in *Xenopus* oocytes

Next, we used CellSpecks to detect channels from experimental data obtained by imaging nAChR activity in *Xenopus* oocytes. As previously shown, TIRFM of Ca^2+^ imaging of membrane regions in oocytes expressing nAChRs revealed numerous transient fluorescence ‘flashes’ (SCCaFTs) in the presence of nicotinic agonists when the membrane is hyperpolarized to increase the driving force for Ca^2+^ influx [15]. Manual analysis of these data revealed a sparse distribution of nAChRs throughout the image field [15]. When we used CellSpecks to detect channels and their events in the same data set, the number of active sites detected was about 3 times larger than the number detected manually. Moreover, CellSpecks required only a few minutes to detect about 850 channels (5178 total events), substantially less than the many hours normally needed for manual inspection. The location map of all the channels and examples of processed fluorescence time-traces from 3 channels given by CellSpecks are shown in Figure 4A and 4B respectively. The ability of CellSpecks to segregate between two nearby channels can be limited in specifics situation in which two channels overlap at two adjacent pixels with both channels displaying long opening time. We have previously estimated that the spatial spread of a SCCaFT generated by the opening of nAChRs measured as the full-width at half-maximal amplitude (FWHM) has a spatial spread of ~ 500 nm [14]. In our experiments, controlled level of expression can easily overcome this problem as the channels imaged so far do not display tendencies to cluster, but a clear random distribution that is normally achieved in relatively low channels overexpression.

**Figure 4.**
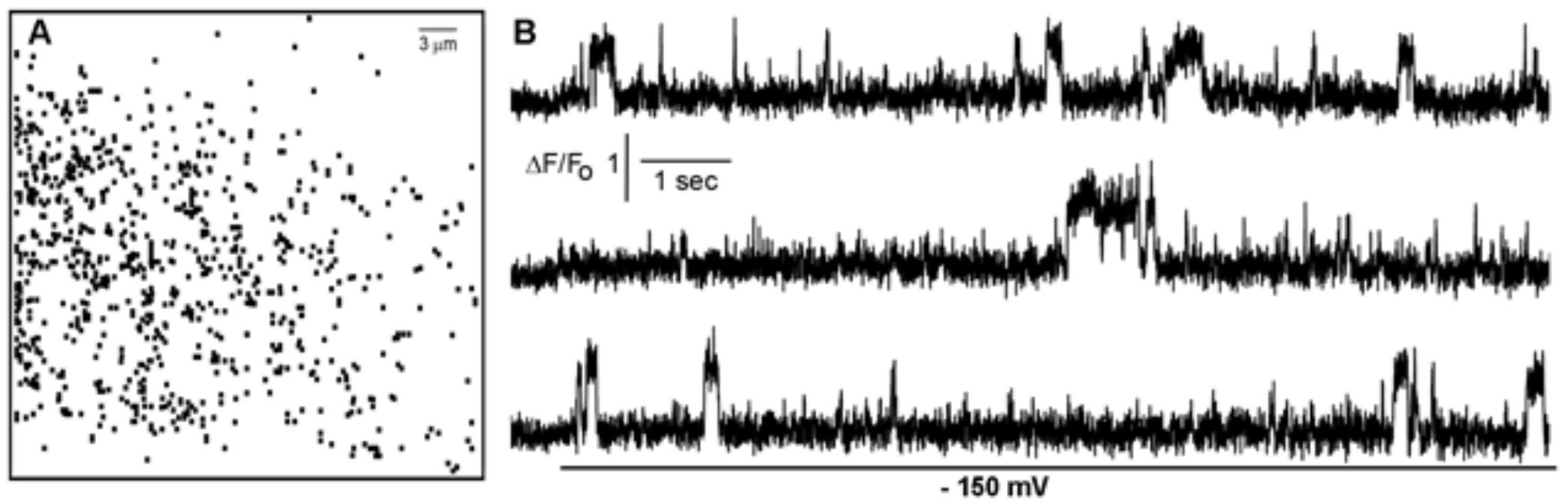
Detection of hundreds of individual nAChR channels by CellSpecks. (A) Channel map showing the locations of 850 nAChRs channels within a 40× 40 μm^2^ membrane patch. The map was generated automatically by CellSpecks after identifying coordinates of all channel sites through a 25 seconds imaging period during which oocyte was polarized to −150 mV in the presence of 1 μM ACh. (B) Example of single-channel recordings (SCCaFTs) resulting from Ca^2+^ influx during channel opening when the membrane was hyperpolarized to −150 mV. Traces are obtained by monitoring fluorescence from regions of interest (1 pixel, corresponding to a 0.33×0.33 μm^2^ of plasma membrane) centered on 3 of the channel locations shown in (A).

The 850 channels detected by CellSpecks included 100% of those detected by hand in experimental datasets. The remaining channels were checked manually in experimental stacks and found to contain at least one centroid uniquely centered at that location, without overlapping nearby channels (occasionally in neighboring pixels) at all. As seen in Figure 5C, the Po distribution indicates that the vast majority of channels detected by CellSpecks are those that have small P_O_ with infrequent and short open events, separated by long close events, affirming the ability of CellSpecks to detect channels with low P_O_ and events with extremely short life-times.

### Automated analysis of channel properties

Analysis of the large amount of information provided by simultaneously analyzing the behavior of a few thousands ion channels is extremely challenging using available data analysis software such as R (GNU), Origin (OriginLab) or software designed for analysis of electrophysiological single channel data.

As described in the “Materials and Methods” section, the statistical analysis module contained in CellSpecks is capable of generating statistical records of the parameters characterizing ion channel behavior. For example, in Figure 5 we show statistical plots describing the distributions of open durations (Figure 5A), close durations (Figure 5B), and the maximum amplitudes (Figure 5D) of the SCCaFTs detected in Figure 4A. The distribution of mean P_O_ of all channels is shown in Figure 5C. The ability of CellSpecks to store the lifetimes and amplitudes of all events for all channels detected (Figure 5) make the more complex analysis such as cross correlation studies between any of the measurable channel parameters a lot easier [13].

**Figure 5.**
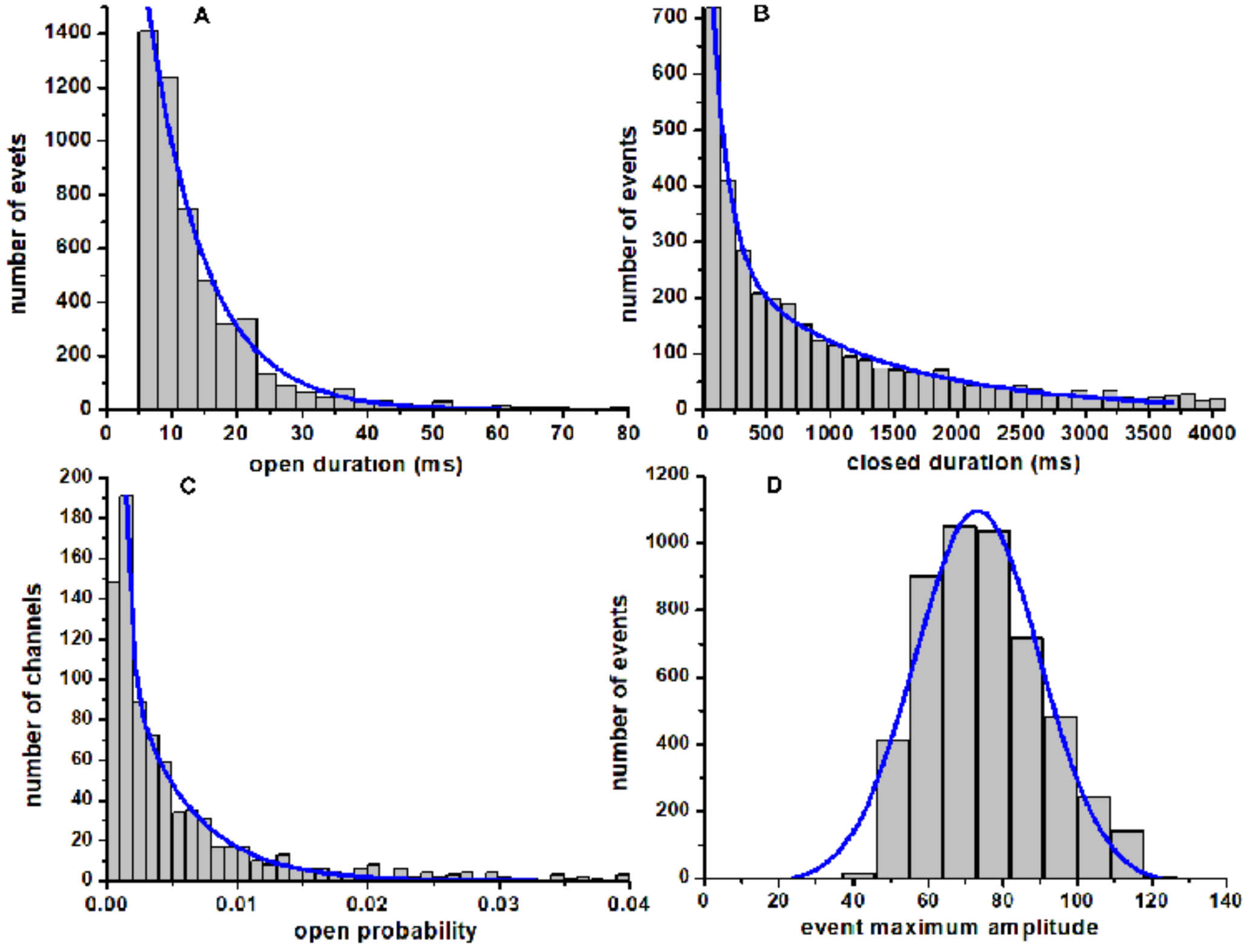
Automated analysis of channel parameters. After locating the active sites (850) in the image field (Fig. 4A), CellSpecks can automatically generate analyses of the fluorescence events (5187) measured during the record, calculating the open times, close times, amplitudes for all events, and mean P_O_ for each corresponding region of interest. These data sets are then used to statistically analyze channel populations. (A) Distribution of the events duration for all of the detected nAChR channels shown in Figure 4A. Data are fitted by single exponential decay (solid line) with time constant 8.6 ms. B) Distribution of the corresponding closed times (intervals between events) fitted by a double exponential decay function (solid line) with time constants of 123.8 ms and 1191.3 ms. C) Distribution of the mean open probabilities calculated for the corresponding channels. Data are fitted by a double exponent decay function with Po1 of 0.00041 and Po2 of 0.0047. (D) Plot displays the maximum events amplitude distribution obtained measuring the peak fluorescence for each detected event. Distribution are fitted by a Gaussian function with a peak amplitude of 73 DF/Fo × 100 (solid line).

### Detection of Aβ42 pore activity

The first version of CellSpecks was developed and used to investigate gating properties of Aβ42 pores in *Xenopus laevis* oocytes [13] allowing the discovery of a large population of Aβ42 pore with very low Po, otherwise overlooked and underestimated by visual inspection. Moreover, CellSpecks is not only capable of correctly and efficiently characterizing the behavior of channels with single permeability level, but can also characterize channels with multiple Ca^2+^ permeability levels. As an example, we applied CellSpecks to TIRFM imaging data of Ca^2+^-permeable plasma membrane pores formed by Aβ42 oligomers. In this case, CellSpecks identified 830 active sites (Figure 6). Distributions for mean open times, mean close times, mean PO, and peak amplitudes for all channels detected are shown in Figure 6A-D. Notice that the distributions shown in Figure 6A and B are the mean values for all channels (one value per channel), which are different than those in Figure 5A and B representing the open and close times for all events (multiple values per channel). Similarly, the amplitude distribution in Figure 6D represents the largest amplitude observed for a given channel, while that in Figure 5D represents the amplitudes of all events detected.

Some of these channels have multiple conductance levels. A representative time trace of such a channel is shown in Figure 6E. An idealized version of the trace using the idealization software TraceSpecks [30] is shown in Figure 6F and G. CellSpecks is also capable of providing “channel chips”, an efficient way of plotting the behavior of a large population of ion channels over time [13]. The channel-chip representation can also provide a way of viewing selected regions to reveal gating characteristics at a finer resolution. Channel chip representation for all 830 Aβ42 pores in a 40μm × 40μm plasma membrane patch of *Xenopus laevis* oocyte is shown in Figure 6H with zoomed in versions in Figure 6I and L. As clear from Figure 6C, majority of these pores have extremely low PO with mixed mean open durations less than 20 ms (Figure 6A) separated by close events that are a few thousand milliseconds long (Figure 6B). The low P_O_ of Aβ42 pores is also confirmed by the channel chip representation, showing significantly longer dark stripes (close events) as compared to brighter dots (open events) (Figure 6H-J).

**Figure 6.**
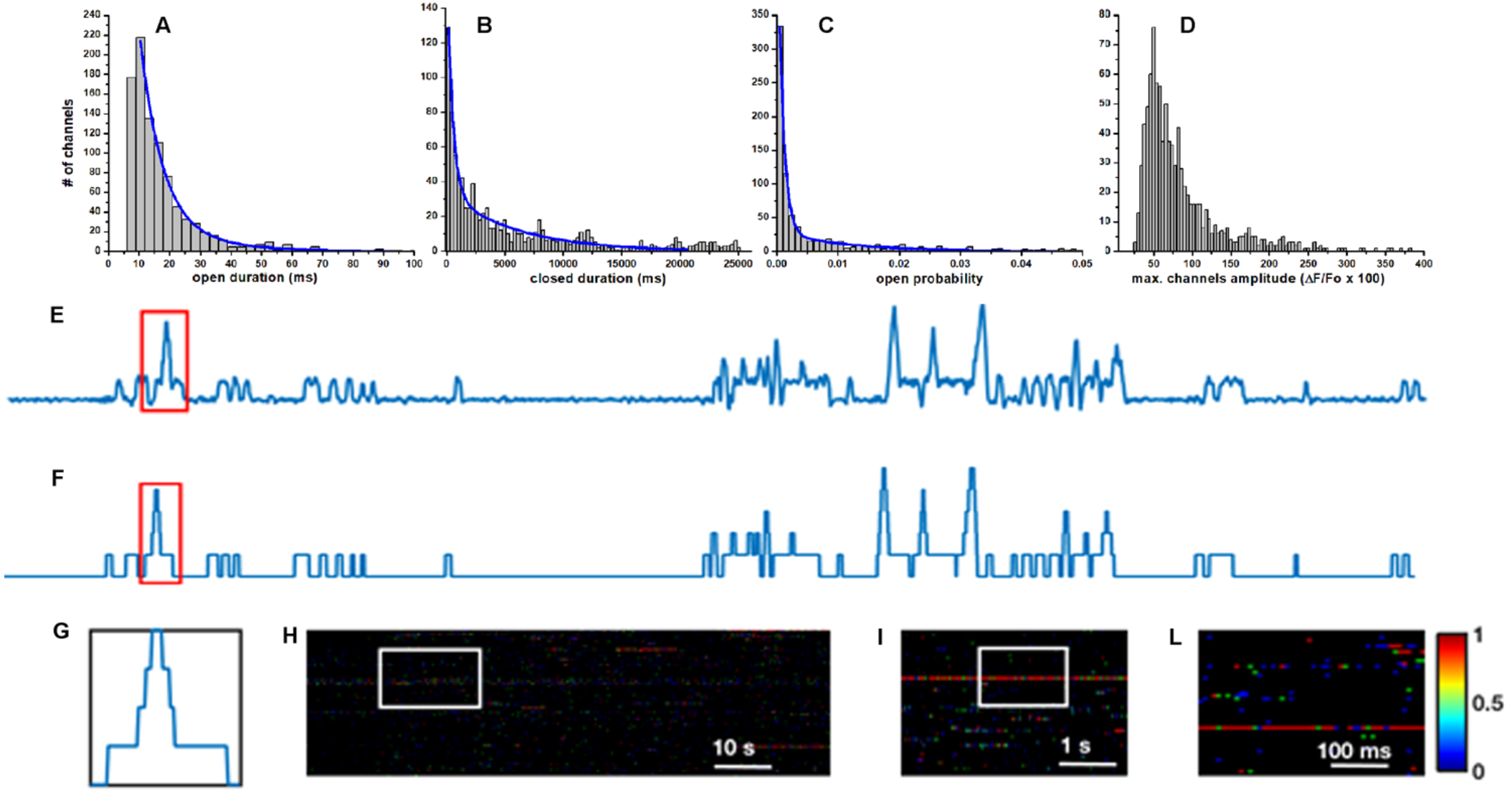
Characterizing the activity of Ca^2+^-permeable ion pores formed by Aβ42 oligomers in a 40× 40μm^2^ plasma membrane patch of a *Xenopus laevis* oocyte, imaged through TIRFM. The stack has 5000 frames recorded at 400 frames/sec. In this example, 1μg/ml of Aβ42 oligomers were applied to the bathing solution and Ca^2+^ influx to the cytoplasm was enhanced by applying a hyperpolarizing potential of −80 mV. (A) Distribution of the channels mean events duration (channels mean open time) for 930 active sites. Double exponential fit (solid curve) yield decay constants of 7.2 and 20.1 ms. (B) Distribution of the mean closed times (times where events were not presents) fitted by a double exponential decay with constants values of 0.473 and 6,034 seconds respectively. Plot in (C), shows corresponding distribution of channels open probability fitted by a double exponential decay function (solid line) with curve yielding values of Po1 0.0084 and o2 0,012. (D) Corresponding distribution of maximum amplitude per channel where variation in amplitude among different channels are evident. (E) Sample fluorescence trace representing the Ca^2+^ influx through a single Aβ42 pore. Multiple conductance levels during individual events are indicated in the fluorescence trace and clearly shown in the idealized trace (F). Panel (G) shows a single event where the pore opens up to four conductance levels. Channel chip representation of the gating of all channels detected in the stack (H) with progressively zoomed in view of shown in (I) and (J).

## DISCUSSION

We developed a method, CellSpecks, that allows rapid and accurate detection, localization and analysis of thousands of functional Ca^2+^-permeable ion channels imaged by TIRFM in intact cells. Although, the program is currently being staged for use in experiments primarily for the detection and analysis of functional Ca^2+^ permeable channels, we believe CellSpecks is equally applicable to the detection and analysis of other channels and molecules studied through fluorescence microscopy [34]. Nevertheless, the development of the software will be ongoing and new features and capabilities will be incorporated as the need arises. Furthermore, although many find the speed of Java to be limiting in such calculation-heavy programs, Java is approaching the speed of C++ for many applications. Improvements to the Java Virtual Machine and compilers will continue the trend toward C++ and Java speed parity [35]. We currently do not see any issue with the performance or platform specificity, however, if needed for speed, interface, or other technical reasons (such as experimentation with CUDA or OpenCL), porting parts or the entirety of the program to another language will be considered in the future.

CellSpecks’ limitations are mostly the result of being built to operate on a particular type of input data. The program only works with signal that deviates from the baseline in a positive direction, and only when the noise is uniformly distributed above and below the baseline. However, this can be easily fixed in the source code (available from the authors upon request) if needed by the experiment. The one known confounding scenario for CellSpecks’ detection algorithm is in differentiating two large nearby events whose fluorescence overlaps both spatially and temporally. Although for ion channels with low PO encountering such scenario is rare, in cases where the P_O_ is high such situation may arise. The solution to this problem will likely come as an event filter that splits into discrete events any event with two or more distinct centroid peaks. Resolving this issue will be coupled with incorporating a feature for tracking the movement of channels (or fluorescence molecules) in the future.

While the specificities discussed above are limitations to the flexibility of the program, they are in many ways what makes this program useful. Using a line-scan detection software like SparkMaster would be ineffective for analyzing 2D image stacks [20]. Similarly, the 2D image stack detection software like GMimPro, developed for Windows operating system, does a very good job at tracking single molecules but is inefficient when a comprehensive statistical analysis for thousands of channel in a single stack is sought due to the significant post-processing required to extract these statistics [19]. Furthermore, the detection of multiple conductance levels and channels with extremely small P_O_ is beyond the scope of GMimPro as it excludes real events when short-lived false events are excluded.

CellSpecks is also advantageous over FLIKA [22] when it comes to the detection of individual channels. CellSpecks focusses on detecting the activity of many simultaneously active single channels in a cell whereas FLIKA is written for analysis of Ca^2+^ puffs resulting from the concerted gating of clustered IP_3_R channels. Apart from differences in the temporal and spatial properties of single channel events and Ca^2+^ puffs these programs were written to detect, there are a few notable differences between the two detection paradigms. CellSpecks computes noise threshold pixel by pixel thus eliminating the need for the user-specified baseline level. FLIKA on the other hand has several thresholds (subtraction of baseline intensity, width of Gaussian filter, frequency thresholds for temporal bandpass filtering etc.) that the user must employ before initializing puff detection. Selection of meaningful thresholds can be a trial and error process. As the choice of thresholds is user-dependent it introduces some arbitrariness in pre-detection processing of the stack and can be time consuming. FLIKA on the other hand is better equipped to deal with changing baselines.

Developing flexible tools for biological discovery drives the need for a very specific set of features. This lends itself to actively developing a piece of software simultaneously to meet these needs. In this case we had two primary requirements for any single piece of software. The first requirement is the ability to accurately detect and localize every channel in the image field. This could not be done manually or by existing packages, especially when the channels have extremely low P_O_ as mentioned above. The second requirement, the ability to export and measure any number of behavioral characteristics, is only solved by being able to design and extract novel measurements from the detection data within the program, a feature not available in any multi-frame analysis software that meets our detection requirements. An additional advantage to home-built software is the ability to easily incorporate statistical routines into the exported material, including parameter correlation and the ability to export extremely large sets of data in virtually any format. CellSpecks allows us to meet both of these requirements, and with a public release provides a tool for researchers solving similar problems.

While manual analysis was clearly missing some small or infrequent events that became apparent on a second close inspection, the sheer number of these legitimate channels, for example, as depicted in Figure 4, expounds the importance of automatic detection and analysis. Without these channels, an analysis of Ca^2+^ flow would be incomplete at best. With this data, we can confidently analyze the population P_O_ and maximum amplitudes. Parameters such as P_O_, the distribution of which is shown, for example in Figure 5C, prove meaningful when compared to the same channel population after the addition of, for instance, an nAChR antagonist. Other parameters, such as the mean open time, mean closed time, maximum amplitude distributions for each channel (e.g. Figure 6A, B, D), and lifetimes and amplitudes of all events detected in the recording (e.g. Figure 6A, B, D) that CellSpecks stores, can be used in much the same manner to infer meaningful relationships. Our ability to determine temporal relationships from the channel chip representation (for example Figure 6H) and spatial relationships from channel location maps (for example Figure 2 left panel) hold endless potential as a beneficiary of CellSpecks’ ability to retain and reuse detection information in memory. Importantly, all this information can be accessed with a few clicks without selecting or adjusting any parameters in the software.

Though the need for yet another of data or image analysis program is not the first conclusion when confronted with a new type of experimental data, the scarcity of signal detection software capable of accepting video data sets and accounting for variable experimental parameters, such as spot size, duration and intensity, drove the desire to have a flexible framework within which current and future experimenters could design and execute sophisticated algorithms and statistical analysis. CellSpecks, the program developed for the purpose of automating localized cell membrane Ca^2+^ event detection is in the short term an effective tool, but more importantly a good basis for future development.

## AUTHOR CONTRIBUTIONS

S I Shah: Performed research, contributed analytic tools, analyzed data, and wrote the paper.

M Smith: Performed research, contributed analytic tools, analyzed data, and wrote the paper.

D Swaminathan: Performed research, contributed analytic tools, analyzed data, and wrote the paper.

I Parker: Designed research and wrote the paper.

G Ullah: Designed research, contributed analytic tools, analyzed data, wrote the paper.

A Demuro: Designed research, contributed analytic tools, analyzed data, wrote the paper.

## ACKNOWLEDGEMENTS

This works was supported by NIH through grants R01 AG053988 (to AD and GU) and R01 GM065830 (to IP).

## Footnotes

To whom correspondence may be addressed. **or**

**Supplementary Information Text 1:** Supplementary methods and flowcharts for the algorithm.

**Supplementary Information Text 2:** Software, User Manual, sample stack file, and sample image sequence.

